# Population selection and sequencing of *C. elegans* wild isolates identifies a region on chromosome III affecting starvation resistance

**DOI:** 10.1101/635417

**Authors:** Amy K. Webster, Anthony Hung, Brad T. Moore, Ryan Guzman, James M. Jordan, Rebecca E. W. Kaplan, Jonathan D. Hibshman, Robyn E. Tanny, Daniel E. Cook, Erik Andersen, L. Ryan Baugh

## Abstract

To understand the genetic basis of complex traits, it is important to be able to efficiently phenotype many genetically distinct individuals. In the nematode *Caenorhabditis elegans*, individuals have been isolated from diverse populations around the globe and whole-genome sequenced. As a result, hundreds of wild strains with known genome sequences can be used for genome-wide association studies (GWAS). However, phenotypic analysis of these strains can be laborious, particularly for quantitative traits requiring multiple measurements per strain. Starvation resistance is likely a fitness-proximal trait for nematodes, and it is related to metabolic disease risk in humans. However, natural variation in *C. elegans* starvation resistance has not been characterized, and precise measurement of the trait is time-intensive. Here, we developed a population selection and sequencing-based approach to phenotype starvation resistance in a pool of 96 wild strains. We used restriction site-associated DNA sequencing (RAD-seq) to infer the frequency of each strain among survivors in a mixed culture over time during starvation. We used manual starvation survival assays to validate the trait data, confirming that strains that increased in frequency over time are starvation-resistant relative to strains that decreased in frequency. These results document natural variation in starvation resistance. Further, we found that variation in starvation resistance is significantly associated with variation at a region on chromosome III. Using a near-isogenic line (NIL), we showed the importance of this genomic interval for starvation resistance. This study demonstrates the feasibility of using population selection and sequencing in an animal model for phenotypic analyses of quantitative traits, reveals natural variation of starvation resistance in *C. elegans*, and identifies a genomic region that contributes to such variation.

## INTRODUCTION

In order to determine the genetic basis of quantitative traits, it is important to first efficiently phenotype many individuals across many genotypes. By using panels of isogenic lines from a model organism of interest, controlled experiments can be performed for particular traits, and biological replicates facilitate repeat measurements of the same genotype. There are a number of model organisms for which panels of isogenic lines have been generated and sequenced, including *Caenorhabditis elegans* (Srivastava et al. 2017; Churchill et al. 2004; Mackay et al. 2012; Noble et al. 2017; Cook et al. 2017; Peter et al. 2018; Weigel 2012). Hundreds of *C. elegans* wild isolates have been collected from around the globe, whole-genome sequenced and made available for use through the *Caenorhabditis elegans* Natural Diversity Resource (CeNDR) (Cook et al. 2017; Hahnel et al. 2018). These lines have the advantage that they represent the natural diversity of the species, rather than being generated through mutagenesis (Jorgensen and Mango 2002) or experimental mutation accumulation (Denver et al. 2009; Denver et al. 2004). As a result, differences in traits across these strains suggest that there is variation in these traits in the wild. In addition, because *C. elegans* reproduces primarily by selfing, these lines are homozygous and isogenic, and they do not suffer from inbreeding depression (Dolgin et al. 2007), which is ideal for quantitative genetics studies.

Progression through the *C. elegans* life cycle depends on food availability. In rich conditions, larvae will develop through four larval stages and become reproductive adults in a couple of days. However, *C. elegans* larvae undergo an alternative developmental program and arrest development in dauer diapause as an alternative to the third larval stage (L3) in response to high population density, limited nutrient availability, or high temperature (Riddle and Albert 1997). Animals can survive dauer arrest for months and resume reproductive development if conditions improve. In a related but distinct phenomenon, L1 larvae that hatch in the complete absence of food arrest development in a state known as “L1 arrest” or “L1 diapause” (Baugh 2013). Larvae can survive starvation during L1 arrest for several weeks, and arrest is reversible upon feeding. Larvae can also arrest development at later stages in response to acute starvation (Schindler, Baugh, and Sherwood 2014), and adults arrest reproduction in response to starvation (Angelo and Van Gilst 2009; Seidel and Kimble 2011). It is relatively easy to produce pure populations of larvae in L1 arrest, facilitating investigation of the starvation response. Notably, *Caenorhabditis* are frequently found in a starved state in the wild (Barriere and Felix 2006), reflecting the importance of starvation resistance to organismal fitness.

The genetic basis of starvation resistance is of particular interest for its relevance to aging, cancer, diabetes, and obesity, as well as its significance as a determinant of evolutionary fitness. L1 arrest and dauer diapause have been termed “ageless” states because length of time spent in arrest has no effect on the duration of adult lifespan after recovery (Klass and Hirsh 1976; Johnson et al. 1984), though larvae in L1 arrest display signs of aging that are actually reversed upon recovery (Roux et al. 2016). In addition, pathways that affect starvation resistance in larvae also affect aging in adults (Baugh 2013). Furthermore, insulin/IGF signaling and AMP-activated protein kinase (AMPK) affect starvation survival in worms and are conserved in mammals, where they are implicated in diabetes and cancer (Li et al. 2015; Malaguarnera and Belfiore 2011). Genetic analysis of *C. elegans* has historically been performed in the genetic background of the commonly used lab strain, N2. Consequently, prior studies have not leveraged the natural variation of starvation resistance among wild isolates to study genetic variants not found in N2 that impact this trait.

Here, we used a panel of 96 whole-genome sequenced wild strains to determine the extent of natural variation in starvation resistance and to identify a genomic interval that contributes to such variation in *C. elegans*. However, independently assaying L1 starvation survival for each strain would have been labor-intensive, and technical variation may have obscured natural variation of the trait. Thus, we sought an approach that would allow us to pool and starve the strains in a single culture, and then use DNA sequencing to infer the relative frequency of survivors over time. In theory, with a single culture and multiplexed measurements, technical variation is reduced, increasing precision of trait measurement. However, precise measurement of survival frequency requires deep sequencing coverage of each strain, making whole-genome sequencing cost-prohibitive. We therefore used RAD-seq as a reduced-representation fractional genome sequencing strategy (Andrews et al. 2016). We used reads including single-nucleotide variants (SNVs) unique to each strain as markers to infer strain frequency, and we used the change in strain frequency over time after recovery from starvation as the quantitative trait value for GWAS. We identified strains that increased or decreased in frequency the most, and we validated that these individual strains are starvation resistant or sensitive using traditional assays. GWAS identified a region on chromosome III significantly associated with increased starvation resistance. We generated near-isogenic lines for the peak on chromosome III, and we determined that this region is causal for differences in starvation resistance. This work demonstrates that population selection and sequencing approaches can be used to phenotype many strains with precision and efficiency. Further, there is natural genetic variation in starvation resistance in *C. elegans*, and a region on chromosome III explains some of that variation.

## MATERIALS AND METHODS

### Strains

Ninety-six *C. elegans* wild isolate strains were used for pooling in the RAD-seq experiment, including AB1, AB4, CB4851, CB4852, CB4853, CB4854, CB4856, CB4857, CB4858, CB4932, CX11262, CX11264, CX11271, CX11285, CX11292, CX11307, CX11314, CX11315, DL200, DL226, DL238, ED3005, ED3011, ED3012, ED3017, ED3040, ED3046, ED3048, ED3049, ED3052, ED3073, ED3077, EG4347, EG4349, EG4724, EG4725, EG4946, JT11398, JU1088, JU1172, JU1200, JU1212, JU1213, JU1242, JU1246, JU1395, JU1400, JU1409, JU1440, JU1491, JU1530, JU1568, JU1580, JU1581, JU1586, JU1652, JU1896, JU258, JU310, JU311, JU323, JU346, JU360, JU363, JU367, JU393, JU394, JU397, JU406, JU440, JU561, JU642, JU751, JU774, JU775, JU778, JU782, JU792, JU830, JU847, KR314, LKC34, LSJ1, MY1, MY10, MY16, MY18, MY23, PB303, PB306, PS2025, PX174, PX179, QX1211, QX1233, WN2002. These strains were obtained from the *Caenorhabditis* Genetics Center (CGC). In addition, Bristol N2 was used in validation experiments, along with QX1430, a modified N2 strain that is genetically compatible with strains that lack *peel-1/zeel-1* (Andersen et al. 2015).

### Worm Culture and Sample Collection for RAD-seq

Ninety-six strains were maintained on *E. coli* OP50-seeded nematode growth medium (NGM) plates at 20°C. Larvae were washed from clean, starved 6 cm plates with S-complete, one plate per strain, and pooled. 20% of this pool was cultured in liquid with 50 mg/mL *E. coli* HB101 in 500 mL total volume of S-complete. After 67 - 72 hours at 20°C and 180 rpm, this culture was hypochlorite treated to obtain an axenic population of embryos, with a yield of 7 - 14 million embryos. Embryos were resuspended at a density of 10 per µL in virgin S-basal (no ethanol or cholesterol). These embryos hatched and entered L1 arrest. For the first biological replicate, two time points were sampled at days 16 and 21 (early time points were sampled, and survivors were separated from dead worms with a sucrose float, but this approach did not work well in later time points, and these samples were abandoned). For the second biological replicate, the culture was sampled at days 1, 7, 14, 21, and 24. For sampling, survival was scored, and approximately 1500 survivors were plated per 10 cm NGM plate with an OP50 lawn (larger volumes of the starvation culture were sampled in later time points as survival decreased; worms were pelleted by centrifuge and plated in 1 mL), with ten plates per time point. Recovery plates were cultured until all OP50 on the plate was consumed. This initially took about four days, but in the later time points additional time was necessary. Starved recovery plates were washed with virgin S-basal. Cultures were spun down at 3000 rpm for 1 minute and excess S-basal was aspirated off. Worms were pipetted into a 1.5 mL tube and flash frozen with liquid nitrogen prior to storage at −80°C. The Qiagen DNAeasy Blood and Tissue column-based kit was used to prepare genomic DNA.

### Restriction site-Associated DNA sequencing (RAD-seq)

RAD-seq was performed as previously described (Davey and Blaxter 2010) in the Duke GCB Sequencing and Genomic Technologies Shared Resource. DNA from each timepoint was digested with *Eco*RI, and adapters were ligated. The ligation products were sheared, and a second adapter was ligated. This second adapter cannot bind to a PCR primer unless its sequence is completed by the amplification starting from a primer bound to the first adapter. This selectively amplified only fragments containing both adapters. The fragments were amplified through PCR and then size selected for 200-500 bp fragments. These fragments were sequenced on an Illumina HiSeq 2500. All sequencing data is archived at NCBI under BioProject number PRJNA533789. The 96 strains used were previously sequenced by RAD-seq, and SNVs were called (Andersen et al. 2012).

### Data Processing

RAD-seq FASTQ files were mapped to version WS210 of the *C. elegans* genome using version 0.7.4 of bwa with default parameters (Li and Durbin 2009). SNVs were called using the command: bcftools call -mv -0z (Li 2011). 29,859,216 to 122,254,549 reads per sample mapped to variants. Nucleotide counts were stored for each of the variants. Counts for all SNVs are available as part of Supplementary File 1.

### Counts to Trend Values for GWAS

Count tables were filtered to only include counts for unique SNVs previously identified (Andersen et al. 2012). The count table for unique SNVs is available as part of Supplementary File 1. Unique SNVs are those for which only one strain out of the 96 has the alternative SNV. For each strain and library/timepoint, the average alternative SNV frequency was calculated, approximating the frequency of each strain. For replicates one and two, linear regressions were calculated with day as the independent variable and the average value for the strain as the dependent variable. The slopes of these regressions indicate whether the representation of a strain tends to increase or decrease over time in culture. The slopes of these linear regressions for replicates one and two were averaged, and these “RAD-seq trait values” were used for GWAS (Supplementary Table 2). The R script to go from unique SNV count values to trait values used for GWAS is available at github.com/amykwebster/RADseq_2019_StarvationResistance

### GWAS using CeNDR

The *Caenorhabditis elegans* Natural Diversity Resource (CeNDR) was used to perform GWAS (Cook et al. 2017) using the EMMA algorithm via the rrBLUP package (Endelman 2011; Kang et al. 2008). The RAD-seq trait values and strain names were used as input for GWAS. The CeNDR version used was 1.2.9, with data release 20180527 and cegwas version 1.01. Version WS263 of the worm genome was used in this data release. Data for strains CB4858 and JU363 were removed to run GWAS because these were considered to be from the same isotype on CeNDR.

### L1 Starvation Survival

Strains of interest were cultured on 10 cm NGM plates seeded with *E. coli* OP50. Gravid adults were hypochlorite treated to obtain axenic embryos. Embryos were re-suspended in virgin S-basal at a density of 1/µL and cultured so they hatch and enter L1 arrest. A 5 mL volume was used in a 16 mm glass test tube with a plastic cap, which was incubated on a tissue culture roller drum at 21-22°C for the duration of the experiment.

Beginning on day 1 and continuing every other day, 100 µL samples were taken from the L1 arrest culture and plated on a 5 cm NGM plate with a spot of OP50 in the center. The L1 larvae were plated around the outside of the spot of OP50. The number of worms plated (T_p_) was counted. Two days later, the number of live worms was counted (T_a_). The percent survival on each day was calculated as total alive divided by total plated (T_a_ / T_p_). For Figure 2A, four biological replicates were scored for seven strains (N2, CB4856, DL200, ED3077, JU1652, JU258, JU561) and three biological replicates were scored for one strain (QX1430). For Figure 4A, four biological replicates were scored for each of the four strains. Logistic curves were fit to each biological replicate and strain in R using the package ‘wormsurvival’ (Kaplan et al. 2015), and the median survival time was calculated. Bartlett’s test was used to test for differences in variances among median survival times across strains. For instances in which there were no significant differences in variance, we pooled the variances across strains for further analysis. We used a one-way ANOVA to first test for differences in average median survival time across strains. If there were differences, we proceeded to perform unpaired t-tests for comparisons of interest. Supplementary Tables 1 and 3 contain summary statistics for starvation survival.

**Figure 1.**
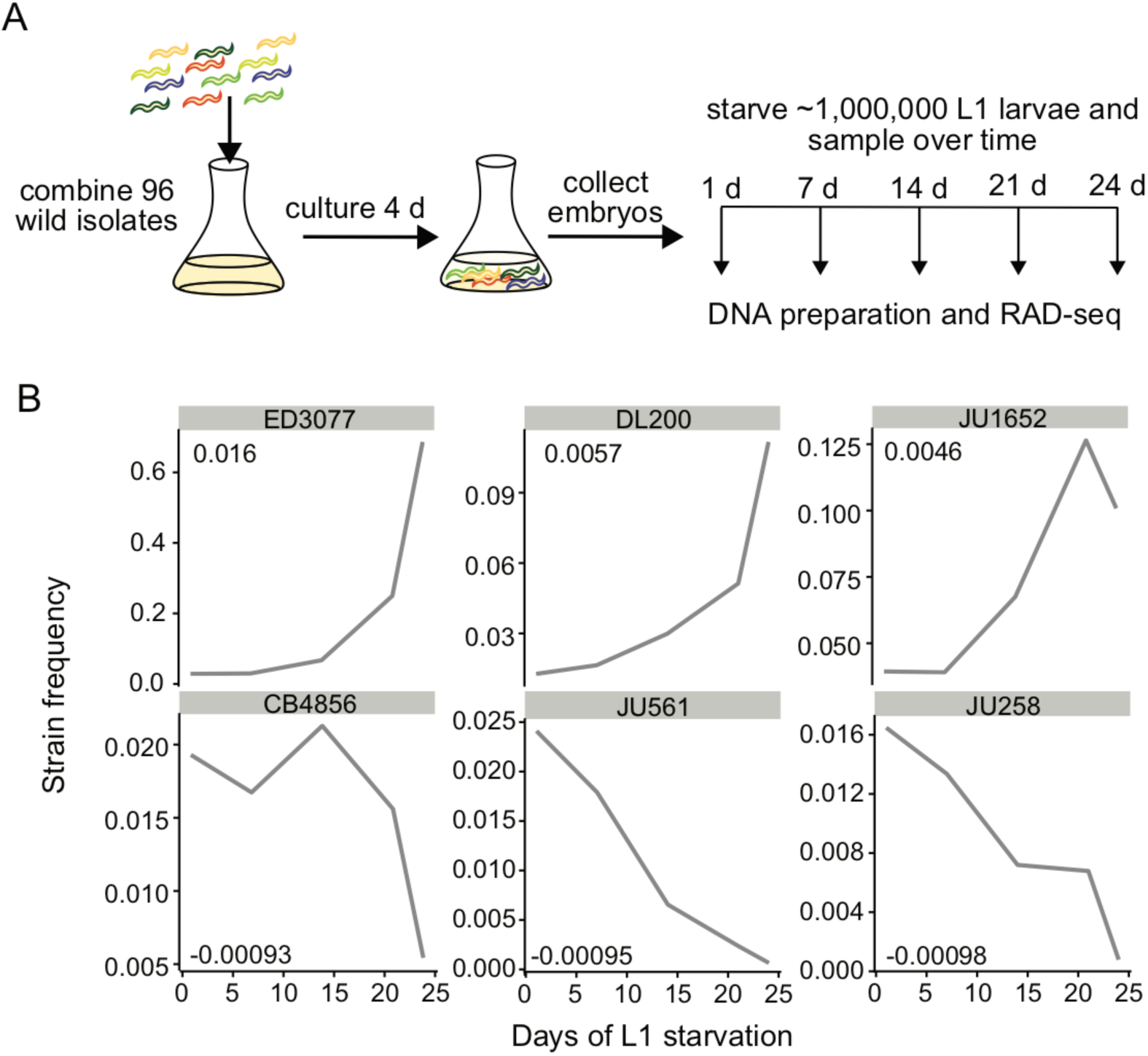
A population selection and sequencing approach to analysis of starvation resistance. A) A schematic of our approach is presented, including pooling of wild strains, co-culture, collection of embryos, establishment of a starvation time series during L1 arrest, and sampling by recovery over time. B) The inferred frequency of strains found to have the highest and lowest RAD-seq trait values is plotted over time during L1 starvation for replicate 2, which included time points spanning early and late starvation. RAD-seq trait values for each strain are included in the corner of each plot. See Supplementary Table 2 for RAD-seq trait values for all strains assayed.

**Figure 2.**
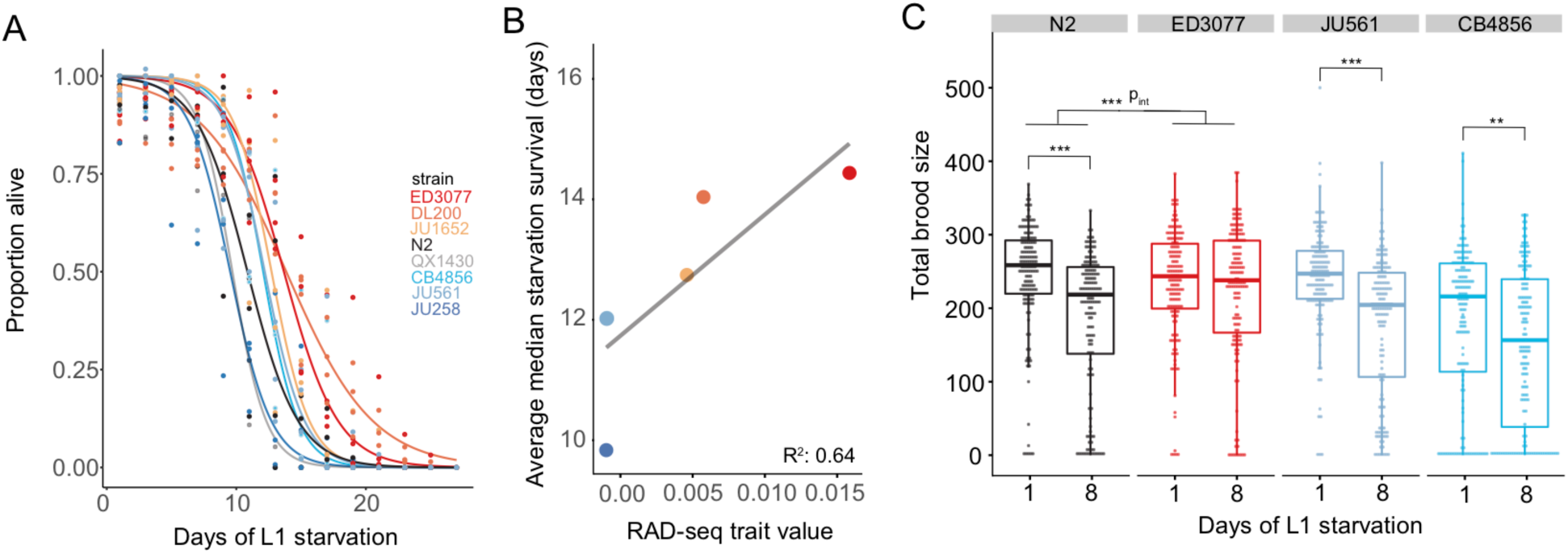
Validation of RAD-seq results with traditional starvation assays. A) L1 starvation-survival curves for strains with the highest and lowest RAD-seq trait values are plotted (see Figure 1B). Starvation survival was assayed for individual strains by manual scoring. Three or four biological replicates were performed, and logistic curves were fit to determine median survival times for each replicate. Summary statistics for starvation survival data are presented in Supplementary Table 1. B) Correlation between RAD-seq trait value and average median starvation survival is plotted. Multiple R^2^ = 0.64, slope of simple linear regression p= 0.055. Note that the point for CB4856 is hidden by the point for JU561. C) Total brood size for N2 (control), a strain found to be starvation resistant (ED3077) and a pair of strains found to be relatively starvation sensitive (JU561 and CB4856) is plotted. Each point indicates the total brood size measured for a single worm. Eighteen individual worms were measured per strain, replicate, and days of L1 arrest. Eight biological replicates were scored for a total of 1,152 worms. Linear mixed-effect models were fit for each strain, with days of L1 arrest as a fixed effect and biological replicate as a random effect. To test for an interaction between strain and days of L1 arrest, data from N2 and ED3077 were fit to a linear mixed-effects model with an interaction term; p_int_ = 0.0001. ***p < 0.001, **p < 0.01.

**Figure 3.**
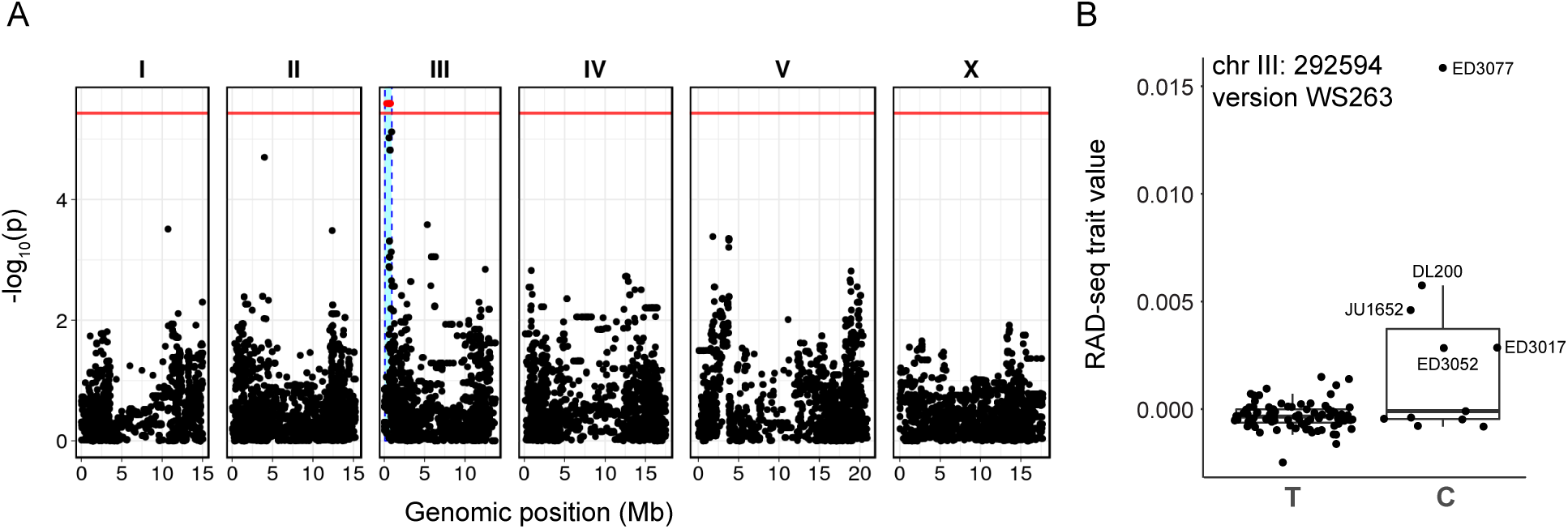
GWAS of RAD-seq trait values identifies a QTL on chromosome III. A) A Manhattan plot for GWAS using RAD-seq trait values (Supplementary Table 2, Supplementary File 1) as input for 96 strains is presented. The horizontal red line indicates the Bonferroni-corrected threshold for statistical significance at a p-value of 0.05, and is 5.43 for this GWAS. A SNV at chromosome III: 292,584 is significantly associated with starvation survival. B) A genotype-by-phenotype plot for the marker SNV most significantly associated with variation in starvation resistance is presented. Each point represents a particular strain, with its genotype at the SNV and its RAD-seq trait value plotted. The most starvation-resistant strains, and a few others, have the alternative allele (C rather than T).

**Figure 4.**
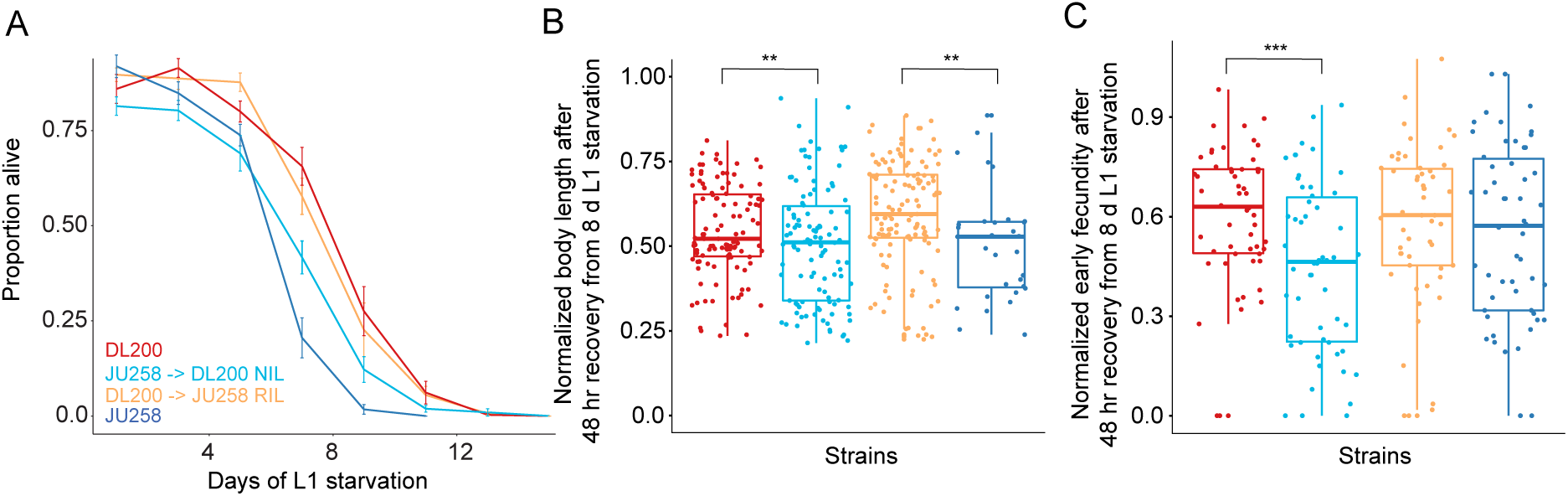
Recombinant strains show that the genomic region on chromosome III associated with variation in starvation resistance affects the trait. A) L1 starvation-survival curves for starvation-resistant strain DL200, starvation-sensitive strain JU258, and newly generated strains DL200 → JU258 RIL and JU258 → DL200 NIL. Four biological replicates were performed, and logistic curves were fit to determine median survival times for each replicate. There were significant differences in median survival across all strains, p = 0.0014, one-way ANOVA. Pairwise t-tests for all four strains were performed. See Results and Supplementary Table 3 for individual corrected p-values and summary statistics. B) Worm length after 48 hours of recovery from eight days of L1 starvation is plotted for the same four strains as in A. Length after eight days of L1 starvation is normalized by length after one day of starvation to isolate the effect of starvation. See Results for individual p-values. C) Brood size on the first day of egg laying (early fecundity; 48 - 72 hr after recovery from starvation by feeding) after recovery from eight days of L1 starvation is plotted. Brood size after eight days of L1 starvation is normalized by brood size after one day of starvation to isolate the effect of starvation on early fecundity. B and C) Each point is an individual worm. Data are pooled from three biological replicates. A linear mixed-effects model was fit to the data using strain as a fixed effect and biological replicate as a random effect. **p < 0.01, ***p < 0.001.

### Total Brood Size

For total brood size (Figure 2C), worms starved as L1 larvae for one day or eight days (as described under ‘L1 Starvation Survival’) were plated on 5 cm NGM plates with OP50 and allowed to recover for 48 hours at 20°C. After 48 hours, worms were singled onto plates with OP50. Every 24 hours, the individual worms were each transferred to a new plate until egg laying ceased (96 hours). 48 hours after eggs were laid, the number of progeny on each plate was scored. The total brood size for each worm was calculated as the total number of progeny laid on all plates for a single worm. 1,152 total individual worms were scored, including eighteen individual worms for each strain (N2, CB4856, JU561, ED3077), time starved (one or eight days), and biological replicate (eight biological replicates). No worms were censored from analysis, including those that died during egg laying or produced no progeny. For statistical analysis, the R package ‘nlme’ was used to fit linear mixed-effects models to the data. For each strain, the number of days of starvation (one or eight) was a fixed effect, and biological replicate was a random effect. To test for interactions between days of starvation and strain, we included both genotype and days of starvation as fixed effects, and included an interaction term in the model. To assign a p-value, we used the “summary” function in R, which performed a t-test on the t-value. For significant p-values, we rejected the null hypothesis that the slope of the linear mixed-effects model for the variable of interest was zero.

### Starvation Recovery and Early Fecundity

L1 larvae were starved for one or eight days (as described under ‘L1 Starvation Survival’). At each time point, larvae were plated on NGM plates with OP50 bacteria and allowed to recover for 48 hours at 20°C. To measure worm body length, after 48 hours of recovery, worms were washed off the recovery plates with virgin S-basal and plated on unseeded 10 cm NGM plates for imaging. Images were taken on a ZeissDiscovery V20 stereomicroscope. Images were analyzed with the WormSizer plugin for FIJI to determine worm length and manually passed or failed (Moore, Jordan, and Baugh 2013). For early fecundity, worms that recovered for 48 hours were singled onto plates with OP50 and allowed to lay embryos for 24 hours and were then removed. After two days, the progeny on the plates were counted. For both body length and early fecundity, measurements after eight days of starvation were normalized by dividing by the average for the same strain after only one day of starvation. Statistics were performed on these normalized values, and a mixed-effects model was implemented using the ‘nlme’ package in R. Strain was a fixed effect and biological replicate was a random effect. For early fecundity, three biological replicates were scored, and eighteen individual worms were scored per replicate per strain. For body length following starvation and recovery, a total of 1,063 worms were included in analysis, including 48 - 82 individual worms measured per replicate and strain following one day of starvation and 8 - 68 individual worms measured for each replicate and strain following eight days of starvation. Lower numbers of individual worms following eight days of starvation were due to the effect of lethality in JU258, as only live worms were scored. Three biological replicates were scored for both assays.

### Generation of Near Isogenic Lines (NILs)

The GWAS led to identification of a significant quantitative trait locus (QTL) located at III:292,594 associated with starvation resistance, and the linked interval consisted of III:86,993-925,945. A starvation-resistant strain, DL200, that has the alternative SNV associated with increased starvation resistance, and a starvation-sensitive strain, JU258, that has the reference SNV at this locus, were chosen to generate NILs. These strains are genetically compatible at *peel-1*/*zeel-1* loci (Seidel, Rockman, and Kruglyak 2008). Two strains were generated: one with the JU258 interval on chromosome III but with the DL200 background (JU258 → DL200, LRB362), and one with the DL200 interval but with the JU258 background (DL200 →JU258, LRB361). DL200 and JU258 worms were crossed to generate heterozygotes, which then self-fertilized to produce F2 progeny. F2 generation worms were singled and genotyped on both sides of the chromosome III peak following egg laying. For this first cross, homozygotes were chosen from both genotypes to continue crosses. From this step onward, separate crosses were performed for each genotype. Worms that were genotyped as homozygous for DL200 were crossed to JU258 males, and F2 progeny that were homozygous at the interval for DL200 were chosen again following genotyping. These were successively crossed to JU258 males, with homozygotes for DL200 chosen in the F2 generation each time. In total, six crosses were performed, such that the DL200 interval should be in a background consisting mostly of the JU258 genome. For the JU258 interval, homozygotes were successively crossed to DL200 males, with JU258 homozygotes at the chromosome III interval chosen each F2 generation. Again, six crosses were performed to generate this strain. After these crosses and genotyping, each strain was selfed for six generations to ensure homozygosity. After generation of NILs, resulting strains and parent strains were whole-genome sequenced at low coverage. All sequencing data is archived on NCBI under BioProject number PRJNA533789. The VCF-kit tool “hmm”, which uses a Hidden Markov Model to infer from which parent a particular genomic region was inherited, was used to validate the sequence of the resulting strains (Cook and Andersen 2017).

### Genotyping for NILs

Primers were used that amplified near each end of the III:86993-925945 interval followed by restriction digest and gel electrophoresis. The primer sequences were generated using the VCF-kit tool “primer snip” for the region of interest (Cook and Andersen 2017). Primers were tested, and working primers for each end of the interval were used for genotyping. PCR was performed using *Taq* DNA polymerase. Both ends of the interval were genotyped to ensure that a recombination event did not break up the interval. The primers amplified in both DL200 and JU258 backgrounds, but the amplified region contains a SNV affecting a restriction site in one genotype but not the other. These primers are: 1) III:83270_F1 (ttggggtactgtagtcggtg), 2) III:83270_R1 (AAGCTCCTTCCACACGTACG), 3) III:729215_F1 (CGTTTGGCACGTACTGAAGC), and 4) III:729215_R3 (AGAACGTCGTAGCCGTCATC). III:83270_F1 and III:83270_R1 function as a primer pair, and *BstUI* is used to digest the PCR products. The PCR product for both DL200 and JU258 is 610 bp. When *BstUI* is used, it cuts the JU258 PCR product into 348 and 262 bp products, but it does not cut the DL200 product. III:729215_F1 and III_729215_R3 function as a primer pair, and *MseI* is used to digest the PCR products. For both JU258 and DL200, the PCR product is 700 bp. Upon digestion, the JU258 product is cut into 118 bp and 582 bp, and the DL200 product is cut into 118, 143, and 439 bp. For both enzymes, digests were performed at 37°C for at least one hour.

The PCR protocol used was:

1. 95°C for 30 seconds
2. 95°C for 15 seconds
3. 55°C for 30 seconds
4. 68°C for 1 minute 20 seconds
5. 68°C for 5 minutes
6. Repeat steps 2 - 4 30×
7. Hold at 15°C

### Data Availability Statement

Raw sequencing data is available through NCBI under BioProject number PRJNA533789. Files to generate trait data for GWAS is available at github.com/amykwebster/RADseq_2019_StarvationResistance, and output from GWAS is available at elegansvariation.org/report/301dcc3f/RADseq_TraitValue. Input and output of GWAS is also available as part of Supplementary File 1, which was deposited to FigShare.

## RESULTS

We pooled 96 wild isolates that had been genotyped by RAD-seq (Andersen et al. 2012) together as larvae, grew them to adulthood in a single large culture, prepared embryos from them, and cultured the embryos in the absence of food so they hatched and entered L1 arrest (Figure 1A). We sampled this starved culture over time by taking aliquots and feeding them in separate recovery cultures. Recovery cultures were allowed to become reproductive and starve out, typically after about one day of egg laying. We isolated DNA from recently starved recovery cultures, including parents and their progeny (see Materials and Methods). Starvation during L1 arrest reduces growth rate and fecundity upon recovery (Jobson et al. 2015), so this approach enriched for haplotypes that retained the greatest fitness following starvation.

We used RAD-seq count data corresponding to unique variants to determine the frequency of each wild strain over time. Focusing on variants that are unique to each strain in the population, we determined the alternative to reference allele ratio from sequencing reads. We used the average of this value across all unique SNVs for each strain as a measure of the frequency of the strain in the population. We then calculated the slope of a linear regression fit to the strain frequencies over time, generating a RAD-seq trait value for each strain (Supplementary Table 2). Theoretically, strains that increased in frequency survived starvation better than other strains and are relatively starvation resistant and *vice versa* for strains that decreased in frequency.

We identified a subset of strains that were particularly starvation resistant or sensitive based on RAD-seq trait values (Figure 1B). To validate these results, we measured both L1 starvation survival and brood size following one or eight days of L1 starvation using traditional assays with individual strains (Figure 2). For L1 starvation survival, we calculated the median survival time for each strain in each replicate after fitting survival curves (Kaplan et al. 2015) (Supplementary Table 1). We found significant differences in the median survival times across all strains (one-way ANOVA, p = 0.0049). Specifically, ED3077 and DL200 increased in frequency over time in the RAD-seq results and had increased starvation survival relative to the standard lab strain, N2 (Figure 2A, p = 0.0045 and p = 0.0098). ED3077 and DL200 also survived starvation longer than JU258, which displayed the largest decrease in frequency in the RAD-seq experiment (Figure 2A, p = 0.00094 and p = 0.0021). We directly compared RAD-seq trait values with median survival times resulting from the standard starvation-survival assay, and these values were positively correlated with an R^2^ of 0.64 (Figure 2B), validating the effectiveness of RAD-seq for identifying starvation-sensitive and resistant strains. There is considerable natural variation in starvation resistance among wild isolates of *C. elegans*, and population selection and sequencing can be used to identify strains that vary for this trait.

To complement the standard L1 starvation survival assay, we also measured brood size following one and eight days of L1 starvation to determine total fecundity (Figure 2C). Fecundity is reduced following eight days of L1 starvation in N2 (Jobson et al. 2015), and early fecundity was integrated into the RAD-seq trait value (see above and Materials and Methods). We hypothesized that starvation-resistant strains would exhibit less of a starvation-dependent reduction in brood size. We counted the total number of progeny from individual worms for the starvation-resistant strain ED3077, in addition to N2 (control), JU561 and CB4856 (both starvation sensitive in the RAD-seq data). For N2, JU561, and CB4856, worms that were starved for eight days produced significantly fewer progeny than worms starved for only one day (p = 1.1 × 10^-8^, p = 3.9 × 10^-11^, and p = 0.0024, respectively), consistent with a starvation-dependent reduction in fecundity. However, ED3077 did not exhibit a significant difference in brood size depending on the duration of starvation (p = 0.30), suggesting that it does not exhibit this starvation-dependent cost. We explicitly tested for an interaction between genotype and duration of starvation in N2 and ED3077 and found a highly significant interaction term (p = 0.0001), suggesting that ED3077 worms produce more progeny than N2 specifically when starved for eight days as L1 larvae. These results collectively suggest that ED3077 is buffered against the reduction in brood size that typically occurs following eight days of L1 starvation. Finally, these results further support the use of RAD-seq on populations to reliably measure differences in life-history traits.

We used the RAD-seq trait values to determine if variation in starvation resistance is associated with genetic variation. We performed GWAS using the *Caenorhabditis elegans* Natural Diversity Resource (CeNDR) (Cook et al. 2017), which implements the EMMA algorithm (Kang et al. 2008) to correct for population structure via the rrBLUP package (Endelman 2011). We used RAD-seq trait values for each strain as input. We found that variation in starvation resistance is significantly associated with genetic variation among the strains used (Figure 3, Supplementary File 1, and elegansvariation.org/report/301dcc3f/RADseq_TraitValue). Specifically, a variant located on chromosome III at base 292,594 in version WS263 of the *C. elegans* genome was significantly associated with increased starvation resistance. By running the EMMA algorithm, CeNDR takes into account linkage disequilibrium, which is prevalent in *C. elegans* (Andersen et al. 2012). As a result, the genomic region associated with the trait includes the variant on chromosome III as well as a linked interval ranging from base pair 86,993 to 925,947. This approximately 900 kilobase (kb) region contains 145 protein-coding genes, including 20 variants (in 16 genes) predicted to have high impact on the function of protein-coding genes (Supplementary File 1).

We used crosses to generate near-isogenic lines (NILs) to determine if the associated region on chromosome III affects starvation resistance in controlled genetic backgrounds. Because a NIL contains the genomic region of interest in a different genetic background, the effect of the region on the trait can be directly tested. We introgressed the region on chromosome III from a starvation-resistant strain into the background of a sensitive strain (indicated as strain X → stain Y), and *vice versa*. We chose strains DL200 and JU258, because DL200 survives significantly longer than JU258 during L1 starvation (Figure 1B, 2A, Supplementary Table 1). DL200 also has the alternative SNV for the peak on chromosome III associated with starvation resistance, and JU258 does not (Figure 3B). In addition, these strains are compatible for mating based on known genetic incompatibilities among wild strains of *C. elegans* (Ben-David, Burga, and Kruglyak 2017; Seidel, Rockman, and Kruglyak 2008). We used a series of repeated backcrosses and genotyping to introgress the region on chromosome III from JU258 into the background of DL200. We performed six backcrosses with genotyping followed by six generations of selfing in constructing the NILs so that any non-selected regions should theoretically represent less than 1% of the genome in the final NIL. We checked the final strains using low-coverage whole genome sequencing. We found that the NIL strain JU258 → DL200 included JU258 sequence on chromosome III from coordinates 1-1,438,286, as expected, as well as a small region on the left end of chromosome I (Supplementary Figure 1). The reciprocal NIL strain DL200 → JU258 contained the DL200 sequence of interest on chromosome III, but it also contained DL200 sequence on each of the other autosomes (Supplementary Figure 1), so we consider this strain a recombinant inbred line (RIL) as opposed to a true NIL. Nonetheless, these strains are sufficient to test the effect of the chromosome III region of interest on starvation resistance.

We measured L1 starvation survival in the strains we constructed (JU258 → DL200 NIL & DL200 → JU258 RIL) as well as the parental strains (DL200 & JU258). We found significant differences in median survival across all strains (p = 0.0014, one-way ANOVA) (Figure 4A, Supplementary Table 3). We found that DL200 survives L1 starvation significantly longer than JU258 (p = 0.00069), validating prior results (Figure 1B, 2A). We were particularly interested in the NIL with the region of interest from JU258 in the background of DL200. Median survival of this strain (JU258 → DL200 NIL) was significantly less than DL200 (p = 0.0039) and DL200 →JU258 RIL (p = 0.013). These results suggest that the DL200 genomic region associated with starvation resistance does in fact promote starvation survival, as survival decreases when this region is swapped for the region from JU258. Median survival of JU258 → DL200 NIL was not significantly different from JU258 (p = 0.31), as if the DL200 background does not contain additional variation that contributes to starvation resistance. However, these data have limited power, and the survival results qualitatively suggest that JU258 → DL200 NIL has increased survival late in starvation compared to JU258, despite no significant effect on median survival (Figure 4A). Conversely, median starvation survival of DL200 → JU258 RIL is significantly greater than JU258 (p = 0.0022) and JU258 → DL200 NIL (p = 0.013), but it is not significantly different than DL200 (p = 0.55). These results further support the conclusion that the DL200 genomic region associated with increased starvation resistance promotes starvation survival because strains carrying it survive longer than strains that do not.

We measured the growth rate and early fecundity of each strain following recovery from one and eight days of L1 starvation to complement analysis of starvation survival. For the RAD-seq-based population sequencing experiment, we collected samples after recovery from starvation, integrating effects on survival with effects on growth rate and early fecundity in the final measurement. We therefore believe that separately measuring growth rate and early fecundity in addition to survival is a more rigorous way to address functional significance of the genomic region of interest on chromosome III. To more closely follow the approach used to generate RAD-seq trait values, we normalized measurements of length (after 48 hours of recovery by feeding) following eight days of L1 arrest by measurements of length following one day of L1 arrest. We expect starvation-sensitive strains to have a larger proportional loss in length following extended starvation than resistant strains. Consistent with our expectation, JU258→ DL200 NIL worms were proportionally shorter than DL200 worms after eight days of L1 arrest (Figure 4B, p = 0.0076). DL200 → JU258 RIL worms were proportionally longer than JU258 worms after eight days of L1 arrest (Figure 4B, p = 0.0045). These results further support the conclusion that the region of interest on chromosome III affects starvation resistance, because worms that are more resistant to L1 starvation grow faster upon recovery.

We also counted the number of progeny produced on the first day of egg laying by single worms after recovery from one and eight days of L1 starvation. We again normalized the results after eight days by those after one day to assess the proportional loss in brood size for each strain. Starvation-sensitive strains are expected to produce a proportionally smaller number of progeny after extended starvation. As expected, JU258 → DL200 NIL produced proportionally fewer progeny following eight days of starvation than DL200 (p = 0.0001), suggesting that the region on chromosome III is important for early fecundity following starvation. Given significant effects on both body length and early fecundity following extended L1 arrest, we conclude that the region of interest on chromosome III contains one or more genetic variants affecting starvation resistance.

## DISCUSSION

We developed a population selection and sequencing-based approach to quantitative genetic analysis of starvation resistance in *C. elegans.* We used RAD-seq to infer strain frequency in a pooled population of 96 sequenced wild isolates during selection for starvation survival. These results together with validation in traditional assays show that substantial natural variation in starvation resistance exists in *C. elegans*, and our data indicate strains that are relatively starvation resistant or sensitive. We also used the trait data for GWAS, identifying a region on chromosome III that is associated with variation in starvation resistance, and we used recombinant strains to determine that this genomic region from the strain DL200 confers starvation resistance. Moreover, this work demonstrates the feasibility of using sequencing of pooled populations of strains for phenotypic analysis of quantitative traits in *C. elegans*. This application of a population selection and sequencing approach to phenotypic analysis in a multicellular model can be readily applied to a variety of other quantitative traits in *C. elegans* as well as other organisms that can be cultured in large numbers. Starvation resistance is likely a fitness-proximal trait in nematodes, and this work establishes a foundation for dissection of this fundamental quantitative trait and identification of genetic variants affecting it.

### Population selection and sequencing-based phenotypic analysis

*C. elegans* are particularly amenable to manipulation in large populations consisting of many strains. Their small size allows for millions of individuals to be present in a single culture, providing power for quantitative analysis. Additionally, inbred strains facilitate accurate phenotype measurements of particular strains by genotyping relatively few markers. We took advantage of RAD-seq to infer strain frequencies in a population under selection. Though convenient and far more affordable than whole-genome sequencing for quantitative analysis, RAD-seq suffers the limitation that only regions of the genome proximal to the *Eco*RI restriction sites are sequenced, the vast majority of which are uninformative for differentiating strains. We found that regressing over all variants captured by RAD-seq was no better at inferring strain frequency than considering only unique SNVs (data not shown), and we chose to focus on unique SNVs. However, the number of unique SNVs per strain varies greatly, with some strains having only one or two unique SNVs (Supplementary File 1). Consequently, sensitivity and precision in measuring strain frequency varied by strain. Future work taking advantage of sequencing methods that target unique SNVs or other strain-specific markers (Mok et al. 2017), or sufficiently deep whole-genome sequencing once affordable, should result in substantially improved trait data from such a population-sequencing approach.

Our experimental approach is intended to capture the effects of early-larval (L1) starvation on organismal fitness. Extended L1 starvation reduces growth rate and fecundity upon recovery (Jobson et al. 2015). Furthermore, L1 starvation survival and the ability to grow and develop upon feeding can be uncoupled (Roux et al. 2016), and effects of L1 starvation on growth rate and fertility upon recovery can be modified without detectable effects on starvation survival (Hibshman, Hung, and Baugh 2016). These observations suggest that the most ecologically and evolutionarily relevant way to assess starvation resistance is by assaying the combined effects of starvation on survival, growth rate, and early fecundity. Our experimental method therefore involved taking samples of a starving L1 culture over time and allowing the worms to recover with food. If worms were dead, they did not recover and contributed little if any DNA to the sample. However, if they survived but were relatively slow to become reproductive adults, or if their fertility was compromised, then their representation was reduced in the DNA that was sequenced. Importantly, because we measured recovery on the first day of starvation, before effects on growth rate or fecundity are detectable (Jobson, Jordan 2015), as well as late time points, strains that simply developed faster or were more fertile in our culture conditions were not considered starvation resistant. That is, starvation-resistant strains were identified as those strains with frequencies that increased over time during starvation. Alternative sampling strategies in future studies could be used to isolate different effects of starvation, potentially identifying genetic variants that affect specific aspects of starvation resistance (*e.g*., mortality, growth rate, or fecundity).

It is important to consider that by sequencing populations we measured relative as opposed to absolute starvation resistance. ED3077 and DL200 increased in frequency over time, suggesting increased starvation resistance relative to other strains. However, strains that decreased in frequency over time are not necessarily particularly starvation sensitive in absolute terms – they are sensitive relative to a majority of other strains in this pooled context. Furthermore, relative survival may depend on the composition of the population. Population density affects L1 starvation survival and gene expression in *C. elegans* (Artyukhin, Schroeder, and Avery 2013; Chan, Rando, and Conine 2018). It is therefore possible that there may be complex effects due to competition for resources or natural variation in production of or sensitivity to pheromones or excreted metabolites. Critically, we confirmed that strains that increased and decreased most in frequency in pooled culture survive starvation relatively longer and shorter, respectively, in individual starvation cultures with manual scoring. In particular, by both RAD-seq and traditional starvation assays, ED3077 and DL200 are starvation resistant relative to other wild isolates and N2, and JU258 is starvation sensitive. Future studies could determine the effects of population composition on population dynamics during starvation, and in particular if there is natural variation in density-dependence of starvation survival.

### Identification of a genomic interval affecting starvation resistance in wild isolates of C. elegans

*C. elegans* displays substantial phenotypic plasticity in response to nutrient availability, similar to other nematodes and many animals in general (Fusco and Minelli 2010; Viney and Diaz 2012). *C. elegans* survives on ephemeral food supplies and thrives with boom-and-bust population dynamics (Schulenburg and Felix 2017). Their ability to adapt to starvation, survive, recover, and reproduce is important for fitness and therefore likely to be under relatively strong selection. As a result, it was unclear at the onset of this study how much natural variation in starvation resistance would be identified. Our results suggest that there is substantial natural variation in starvation resistance. In addition to variation at the strain level, one might imagine that the trait is so polygenic or regulated by many interacting loci that genetic variants that influence it may be numerous and of small effect or not able to be readily identified. Efforts to map starvation resistance in *Drosophila* have identified multiple genomic loci, demonstrating the polygenicity of the trait in another model system (Everman et al. 2019). Our identification of a genomic interval affecting starvation resistance suggests that such variants exist in *C. elegans*. However, this interval explains ∼2% of the variation in starvation resistance, and additional studies will likely identify additional loci to further explain this presumably polygenic trait.

We took advantage of the extensive genetic and genomic resources available in *C. elegans* to perform a statistical genetic analysis of starvation resistance. Many *C. elegans* wild isolates have had their genomes sequenced with variation called (Cook et al. 2017). Population genetic analyses have been performed, and relatedness among individual strains is known. We took advantage of these advances and used CeNDR to perform GWAS to determine if any regions of the genome are associated with starvation resistance using RAD-seq trait values from 96 strains. A 900 kb region on the left side of chromosome III was found to be associated with the trait. *C. elegans* is known to exhibit extensive linkage disequilibrium, which can lead to relatively large haplotypes, spanning between dozens of kb to greater than one megabase (Mb) (Andersen et al. 2012). However, one advantage of using *C. elegans* is that inbred lines can be relatively easily generated. To test functional significance of this genomic region, we generated a NIL that contained the region of interest from a starvation-sensitive strain, JU258, introgressed into the background of a starvation-resistant strain, DL200. Despite containing ∼98% of the genome of DL200, this NIL survived L1 starvation significantly less than DL200, and survival did not differ significantly from the starvation-sensitive parental strain, JU258. This suggests that this region is important for conferring starvation resistance in DL200. Consistent with this result, the survival of a RIL that contained the chromosome III region from DL200 but also contained a majority of the JU258 genetic background was not statistically different from DL200. Analysis of growth rate and early fecundity following eight days of starvation also support the conclusion that this genomic region confers starvation resistance. Though we showed this region on the left side of chromosome III is important for starvation resistance, it is unknown whether there is one particular variant that confers resistance or whether there is polygenicity for the trait within the region.

The genetic basis of natural variation in starvation resistance in *C. elegans* remains to be determined. The genomic region on chromosome III explained ∼2% of the total variation in the RAD-seq trait values. Although follow-up phenotypic analysis demonstrated that this region explained a significant proportion of the variation between the JU258 and DL200 strains in particular, we found additional variation in starvation resistance across all strains for which causal genomic intervals have yet to be identified. New strains are being catalogued and sequenced regularly, adding to the genetic diversity of *C. elegans* and providing additional leverage for analysis of starvation resistance and other traits. In future studies, choosing the most diverse subset of strains for the species should further harness the power of population selection and sequencing without requiring large numbers strains, though more strains could also be used. Additional annotation of genetic variation beyond SNVs, and *de novo* assembly of particularly divergent strains, will also increase power of the approach. Population selection and sequencing-based approaches together with ever richer genomic resources should provide the power to dissect a variety of complex quantitative traits.

## ACKNOWLEDGEMENTS

Some strains were provided by the CGC, which is funded by NIH Office of Research Infrastructure Programs (P40 OD010440). A.K.W. is supported by the National Science Foundation Graduate Research Fellowship. This work was funded by the National Institutes of Health (R01GM117408, L.R.B.; R21HD081177, L.R.B; and a training grant for the University Program in Genetics and Genomics 5T32GM007754-38). We thank Olivier Fedrigo and the Duke GCB Sequencing and Genomic Technologies Shared Resource for preparing and sequencing the RAD-seq libraries.

---

**Supplementary Figure 1.**
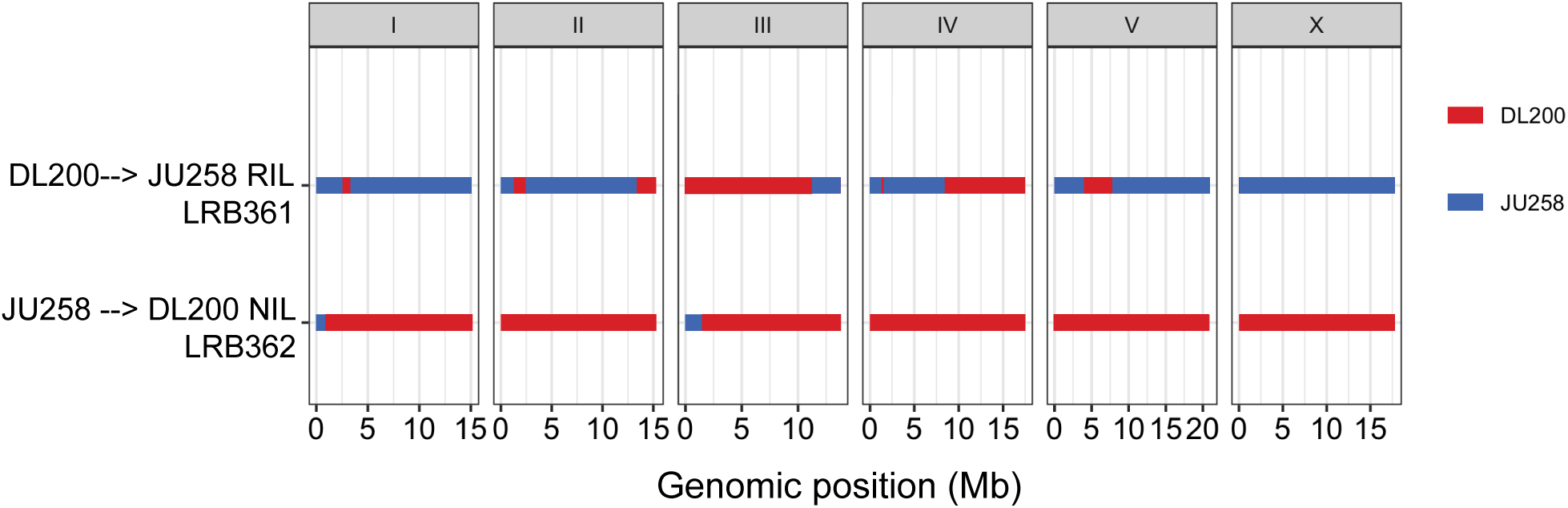
DNA sequencing of near-isogenic lines. The JU258 → DL200 NIL and DL200 → JU258 RIL were sequenced to determine which regions of the genome came from which parental background. Regions of the genome that originated from DL200 and JU258 are color-coded in red and blue, respectively.

**Supplementary Table 1.** Starvation survival summary statistics for Figure 2A.

| Strain | Average median survival (days) | Standard deviation (days) | Number of reps | t-test vs. CB4856 | t-test vs. DL200 | t-test vs. ED3077 | t-test vs. JU1652 | t-test vs. JU258 | t-test vs. JU561 | t-test vs. N2 |
| --- | --- | --- | --- | --- | --- | --- | --- | --- | --- | --- |
| **CB4856** | 12.03 | 1.89 | 4 | n/a | 0.11 | 0.059 | 0.56 | 0.083 | 0.99 | 0.26 |
| **DL200** | 14.04 | 1.36 | 4 | 0.11 | n/a | 0.74 | 0.30 | 0.0021 | 0.11 | 0.0098 |
| **ED3077** | 14.44 | 2.30 | 4 | 0.059 | 0.74 | n/a | 0.18 | 0.00094 | 0.058 | 0.0045 |
| **JU1652** | 12.74 | 1.69 | 4 | 0.56 | 0.30 | 0.18 | n/a | 0.025 | 0.56 | 0.094 |
| **JU258** | 9.84 | 1.82 | 4 | 0.083 | 0.0021 | 0.00094 | 0.025 | n/a | 0.084 | 0.52 |
| **JU561** | 12.02 | 1.48 | 4 | 0.99 | 0.11 | 0.058 | 0.56 | 0.084 | n/a | 0.26 |
| **N2** | 10.62 | 1.53 | 4 | 0.26 | 0.0098 | 0.0045 | 0.094 | 0.52 | 0.26 | n/a |
| **QX1430** | 9.77 | 1.31 | 3 | 0.098 | 0.0035 | 0.0017 | 0.033 | 0.96 | 0.10 | 0.52 |

**Supplementary Table 2.** RAD-seq trait data used for GWAS

| Strain | Trait value | Strain | Trait value | Strain | Trait value |
| --- | --- | --- | --- | --- | --- |
| AB1 | -0.001089558 | EG4946 | -0.001189646 | JU751 | -0.000343356 |
| AB4 | -0.000735581 | JT11398 | -0.000251866 | JU774 | 0.000156211 |
| CB4851 | -1.28E-05 | JU1088 | -0.000542719 | JU775 | -0.00013356 |
| CB4852 | -9.77E-05 | JU1172 | -0.000573628 | JU778 | -0.000466356 |
| CB4853 | -0.000104916 | JU1200 | 0.000643843 | JU782 | -0.000251516 |
| CB4854 | -0.000834588 | JU1212 | -0.000501071 | JU792 | -0.000126172 |
| CB4856 | -0.000926858 | JU1213 | 0.00010362 | JU830 | -0.000421824 |
| CB4857 | -0.000486995 | JU1242 | -0.000508404 | JU847 | 0.000694595 |
| CB4858 | -0.000279677 | JU1246 | -0.000352744 | KR314 | -0.000739883 |
| CB4932 | 8.25E-05 | JU1395 | -0.000812441 | LKC34 | 0.000101218 |
| CX11262 | -0.000776708 | JU1400 | -9.72E-05 | LSJ1 | 0.000228846 |
| CX11264 | -0.001023059 | JU1409 | -0.000496307 | MY1 | -0.000355746 |
| CX11271 | -0.000285448 | JU1440 | -0.00046671 | MY10 | -0.000267622 |
| CX11285 | -0.000454118 | JU1491 | -0.00018339 | MY16 | -0.001084975 |
| CX11292 | 0.000365799 | JU1530 | -0.000394825 | MY18 | 0.000116325 |
| CX11307 | -0.000483986 | JU1568 | -9.76E-06 | MY23 | -0.000536114 |
| CX11314 | -0.000528437 | JU1580 | 0.000636567 | PB303 | -0.001618959 |
| CX11315 | 0.001110572 | JU1581 | -0.001178581 | PB306 | -0.000679825 |
| DL200 | 0.005747001 | JU1586 | -0.000505315 | PS2025 | -0.002479676 |
| DL226 | -0.000764146 | JU1652 | 0.004598108 | PX174 | -0.000170046 |
| DL238 | -0.000804199 | JU1896 | 0.001495203 | PX179 | -0.000586876 |
| ED3005 | 0.001394962 | JU258 | -0.000978538 | QX1211 | -0.000186515 |
| ED3011 | -0.000241451 | JU310 | -0.000783759 | QX1233 | -0.000967571 |
| ED3012 | -0.000347688 | JU311 | -0.000460506 | WN2002 | -0.000671692 |
| ED3017 | 0.002837005 | JU323 | -0.000451522 |  |  |
| ED3040 | -2.54E-05 | JU346 | -0.000320575 |  |  |
| ED3046 | -0.000153265 | JU360 | -0.000889179 |  |  |
| ED3048 | 0.000259088 | JU363 | 0.000173825 |  |  |
| ED3049 | -0.000428888 | JU367 | -0.000317346 |  |  |
| ED3052 | 0.002854565 | JU393 | -0.00097683 |  |  |
| ED3073 | -0.000480596 | JU394 | 8.55E-05 |  |  |
| ED3077 | 0.015852705 | JU397 | -1.14E-05 |  |  |
| EG4347 | 0.000265527 | JU406 | 7.70E-05 |  |  |
| EG4349 | 0.000949259 | JU440 | -0.000588908 |  |  |
| EG4724 | -0.000240064 | JU561 | -0.000947311 |  |  |
| EG4725 | 0.000711111 | JU642 | -0.000175764 |  |  |

**Supplementary Table 3.** Starvation survival summary statistics for Figure 4A.

| Strain | Average median survival (days) | Standard deviation (days) | Number of replicates | t-test vs. DL200 → JU258 RIL | t-test vs. DL200 | t-test vs. JU258 → DL200 NIL | t-test vs. JU258 |
| --- | --- | --- | --- | --- | --- | --- | --- |
| DL200 → JU258 RIL (LRB361) | 7.47 | 0.72 | 5 | n/a | 0.55 | 0.013 | 0.0022 |
| DL200 | 7.72 | 0.77 | 5 | 0.55 | n/a | 0.0039 | 0.00069 |
| JU258 → DL200 NIL (LRB362) | 6.32 | 0.65 | 5 | 0.013 | 0.0039 | n/a | 0.31 |
| JU258 | 5.89 | 0.23 | 4 | 0.0022 | 0.00069 | 0.31 | n/a |

Supplementary File 1. RAD-seq count data for SNVs, trait values for 96 strains, and GWAS output (Excel file). This file contains a readme sheet describing the contents of the file as well as separate sheets for RAD-seq data, including counts of reads mapping to SNVs, unique SNVs for each strain, and trait data. It also includes sheets with the output of GWAS from CeNDR, including a mapping summary, interval summary, and interval variants.

## REFERENCES

Andersen, E. C., J. P. Gerke, J. A. Shapiro, J. R. Crissman, R. Ghosh, J. S. Bloom, M. A. Felix, and L. Kruglyak. 2012. ‘Chromosome-scale selective sweeps shape Caenorhabditis elegans genomic diversity’, Nat Genet, 44: 285-90.

Andersen, E. C., T. C. Shimko, J. R. Crissman, R. Ghosh, J. S. Bloom, H. S. Seidel, J. P. Gerke, and L. Kruglyak. 2015. ‘A Powerful New Quantitative Genetics Platform, Combining Caenorhabditis elegans High-Throughput Fitness Assays with a Large Collection of Recombinant Strains’, G3 (Bethesda), 5: 911-20.

Andrews, K. R., J. M. Good, M. R. Miller, G. Luikart, and P. A. Hohenlohe. 2016. ‘Harnessing the power of RADseq for ecological and evolutionary genomics’, Nat Rev Genet, 17: 81-92.

Angelo, G., and M. R. Van Gilst. 2009. ‘Starvation protects germline stem cells and extends reproductive longevity in C. elegans’, Science, 326: 954–8.

Artyukhin, A. B., F. C. Schroeder, and L. Avery. 2013. ‘Density dependence in Caenorhabditis larval starvation’, Sci Rep, 3: 2777.

Barriere, A., and M. A. Felix. 2006. ‘Isolation of C. elegans and related nematodes’, WormBook: 1–9.

Baugh, L. R. 2013. ‘To grow or not to grow: nutritional control of development during Caenorhabditis elegans L1 arrest’, Genetics, 194: 539–55.

Ben-David, E., A. Burga, and L. Kruglyak. 2017. ‘A maternal-effect selfish genetic element in Caenorhabditis elegans’, Science, 356: 1051–55.

Chan, I. L., O. J. Rando, and C. C. Conine. 2018. ‘Effects of Larval Density on Gene Regulation in Caenorhabditis elegans During Routine L1 Synchronization’, G3 (Bethesda), 8: 1787–93.

Churchill, G. A., D. C. Airey, H. Allayee, J. M. Angel, A. D. Attie, J. Beatty, W. D. Beavis, J. K. Belknap, B. Bennett, W. Berrettini, A. Bleich, M. Bogue, K. W. Broman, K. J. Buck, E. Buckler, M. Burmeister, E. J. Chesler, J. M. Cheverud, S. Clapcote, M. N. Cook, R. D. Cox, J. C. Crabbe, W. E. Crusio, A. Darvasi, C. F. Deschepper, R. W. Doerge, C. R. Farber, J. Forejt, D. Gaile, S. J. Garlow, H. Geiger, H. Gershenfeld, T. Gordon, J. Gu, W. Gu, G. de Haan, N. L. Hayes, C. Heller, H. Himmelbauer, R. Hitzemann, K. Hunter, H. C. Hsu, F. A. Iraqi, B. Ivandic, H. J. Jacob, R. C. Jansen, K. J. Jepsen, D. K. Johnson, T. E. Johnson, G. Kempermann, C. Kendziorski, M. Kotb, R. F. Kooy, B. Llamas, F. Lammert, J. M. Lassalle, P. R. Lowenstein, L. Lu, A. Lusis, K. F. Manly, R. Marcucio, D. Matthews, J. F. Medrano, D. R. Miller, G. Mittleman, B. A. Mock, J. S. Mogil, X. Montagutelli, G. Morahan, D. G. Morris, R. Mott, J. H. Nadeau, H. Nagase, R. S. Nowakowski, B. F. O’Hara, A. V. Osadchuk, G. P. Page, B. Paigen, K. Paigen, A. A. Palmer, H. J. Pan, L. Peltonen-Palotie, J. Peirce, D. Pomp, M. Pravenec, D. R. Prows, Z. Qi, R. H. Reeves, J. Roder, G. D. Rosen, E. E. Schadt, L. C. Schalkwyk, Z. Seltzer, K. Shimomura, S. Shou, M. J. Sillanpaa, L. D. Siracusa, H. W. Snoeck, J. L. Spearow, K. Svenson, L. M. Tarantino, D. Threadgill, L. A. Toth, W. Valdar, F. P. de Villena, C. Warden, S. Whatley, R. W. Williams, T. Wiltshire, N. Yi, D. Zhang, M. Zhang, F. Zou, and Consortium Complex Trait. 2004. ‘The Collaborative Cross, a community resource for the genetic analysis of complex traits’, Nat Genet, 36: 1133–7.

Cook, D. E., and E. C. Andersen. 2017. ‘VCF-kit: assorted utilities for the variant call format’, Bioinformatics, 33: 1581–82.

Cook, D. E., S. Zdraljevic, J. P. Roberts, and E. C. Andersen. 2017. ‘CeNDR, the Caenorhabditis elegans natural diversity resource’, Nucleic Acids Res, 45: D650–D57.

Davey, J. W., and M. L. Blaxter. 2010. ‘RADSeq: next-generation population genetics’, Brief Funct Genomics, 9: 416–23.

Denver, D. R., P. C. Dolan, L. J. Wilhelm, W. Sung, J. I. Lucas-Lledo, D. K. Howe, S. C. Lewis, K. Okamoto, W. K. Thomas, M. Lynch, and C. F. Baer. 2009. ‘A genome-wide view of Caenorhabditis elegans base-substitution mutation processes’, Proc Natl Acad Sci U S A, 106: 16310–4.

Denver, D. R., K. Morris, M. Lynch, and W. K. Thomas. 2004. ‘High mutation rate and predominance of insertions in the Caenorhabditis elegans nuclear genome’, Nature, 430: 679–82.

Dolgin, E. S., B. Charlesworth, S. E. Baird, and A. D. Cutter. 2007. ‘Inbreeding and outbreeding depression in Caenorhabditis nematodes’, Evolution, 61: 1339–52.

Endelman, J. B. 2011. ‘Ridge Regression and Other Kernels for Genomic Selection with R Package rrBLUP’, Plant Genome, 4: 250–55.

Everman, E. R., C. L. McNeil, J. L. Hackett, C. L. Bain, and S. J. Macdonald. 2019. ‘Dissection of Complex, Fitness-Related Traits in Multiple Drosophila Mapping Populations Offers Insight into the Genetic Control of Stress Resistance’, Genetics, 211: 1449–67.

Fusco, G., and A. Minelli. 2010. ‘Phenotypic plasticity in development and evolution: facts and concepts. Introduction’, Philos Trans R Soc Lond B Biol Sci, 365: 547–56.

Hahnel, S. R., S. Zdraljevic, B. C. Rodriguez, Y. Zhao, P. T. McGrath, and E. C. Andersen. 2018. ‘Extreme allelic heterogeneity at a Caenorhabditis elegans beta-tubulin locus explains natural resistance to benzimidazoles’, PLoS Pathog, 14: e1007226.

Hibshman, J. D., A. Hung, and L. R. Baugh. 2016. ‘Maternal Diet and Insulin-Like Signaling Control Intergenerational Plasticity of Progeny Size and Starvation Resistance’, PLoS Genet, 12: e1006396.

Jobson, M. A., J. M. Jordan, M. A. Sandrof, J. D. Hibshman, A. L. Lennox, and L. R. Baugh. 2015. ‘Transgenerational Effects of Early Life Starvation on Growth, Reproduction, and Stress Resistance in Caenorhabditis elegans’, Genetics, 201: 201–12.

Johnson, T. E., D. H. Mitchell, S. Kline, R. Kemal, and J. Foy. 1984. ‘Arresting development arrests aging in the nematode Caenorhabditis elegans’, Mech Ageing Dev, 28: 23–40.

Jorgensen, E. M., and S. E. Mango. 2002. ‘The art and design of genetic screens: caenorhabditis elegans’, Nat Rev Genet, 3: 356–69.

Kang, H. M., N. A. Zaitlen, C. M. Wade, A. Kirby, D. Heckerman, M. J. Daly, and E. Eskin. 2008. ‘Efficient control of population structure in model organism association mapping’, Genetics, 178: 1709–23.

Kaplan, R. E., Y. Chen, B. T. Moore, J. M. Jordan, C. S. Maxwell, A. J. Schindler, and L. R. Baugh. 2015. ‘dbl-1/TGF-beta and daf-12/NHR Signaling Mediate Cell-Nonautonomous Effects of daf-16/FOXO on Starvation-Induced Developmental Arrest’, PLoS Genet, 11: e1005731.

Klass, M., and D. Hirsh. 1976. ‘Non-ageing developmental variant of Caenorhabditis elegans’, Nature, 260: 523–5.

Li, H. 2011. ‘A statistical framework for SNP calling, mutation discovery, association mapping and population genetical parameter estimation from sequencing data’, Bioinformatics, 27: 2987–93.

Li, H., and R. Durbin. 2009. ‘Fast and accurate short read alignment with Burrows-Wheeler transform’, Bioinformatics, 25: 1754–60.

Li, W., S. M. Saud, M. R. Young, G. Chen, and B. Hua. 2015. ‘Targeting AMPK for cancer prevention and treatment’, Oncotarget, 6: 7365–78.

Mackay, T. F., S. Richards, E. A. Stone, A. Barbadilla, J. F. Ayroles, D. Zhu, S. Casillas, Y. Han, M. M. Magwire, J. M. Cridland, M. F. Richardson, R. R. Anholt, M. Barron, C. Bess, K. P. Blankenburg, M. A. Carbone, D. Castellano, L. Chaboub, L. Duncan, Z. Harris, M. Javaid, J. C. Jayaseelan, S. N. Jhangiani, K. W. Jordan, F. Lara, F. Lawrence, S. L. Lee, P. Librado, R. S. Linheiro, R. F. Lyman, A. J. Mackey, M. Munidasa, D. M. Muzny, L. Nazareth, I. Newsham, L. Perales, L. L. Pu, C. Qu, M. Ramia, J. G. Reid, S. M. Rollmann, J. Rozas, N. Saada, L. Turlapati, K. C. Worley, Y. Q. Wu, A. Yamamoto, Y. Zhu, C. M. Bergman, K. R. Thornton, D. Mittelman, and R. A. Gibbs. 2012. ‘The Drosophila melanogaster Genetic Reference Panel’, Nature, 482: 173–8.

Malaguarnera, R., and A. Belfiore. 2011. ‘The insulin receptor: a new target for cancer therapy’, Front Endocrinol (Lausanne), 2: 93.

Mok, C. A., V. Au, O. A. Thompson, M. L. Edgley, L. Gevirtzman, J. Yochem, J. Lowry, N. Memar, M. R. Wallenfang, D. Rasoloson, B. Bowerman, R. Schnabel, G. Seydoux, D. G. Moerman, and R. H. Waterston. 2017. ‘MIP-MAP: High-Throughput Mapping of Caenorhabditis elegans Temperature-Sensitive Mutants via Molecular Inversion Probes’, Genetics, 207: 447–63.

Moore, B. T., J. M. Jordan, and L. R. Baugh. 2013. ‘WormSizer: high-throughput analysis of nematode size and shape’, PLoS One, 8: e57142.

Noble, L. M., I. Chelo, T. Guzella, B. Afonso, D. D. Riccardi, P. Ammerman, A. Dayarian, S. Carvalho, A. Crist, A. Pino-Querido, B. Shraiman, M. V. Rockman, and H. Teotonio. 2017. ‘Polygenicity and Epistasis Underlie Fitness-Proximal Traits in the Caenorhabditis elegans Multiparental Experimental Evolution (CeMEE) Panel’, Genetics, 207: 1663–85.

Peter, J., M. De Chiara, A. Friedrich, J. X. Yue, D. Pflieger, A. Bergstrom, A. Sigwalt, B. Barre, K. Freel, A. Llored, C. Cruaud, K. Labadie, J. M. Aury, B. Istace, K. Lebrigand, P. Barbry, S. Engelen, A. Lemainque, P. Wincker, G. Liti, and J. Schacherer. 2018. ‘Genome evolution across 1,011 Saccharomyces cerevisiae isolates’, Nature, 556: 339–44.

Riddle, D. L., and P. S. Albert. 1997. ‘Genetic and Environmental Regulation of Dauer Larva Development.’ in D. L. Riddle, T. Blumenthal, B. J. Meyer and J. R. Priess (eds.), C. elegans II(Cold Spring Harbor (NY)).

Roux, A. E., K. Langhans, W. Huynh, and C. Kenyon. 2016. ‘Reversible Age-Related Phenotypes Induced during Larval Quiescence in C. elegans’, Cell Metab, 23: 1113–26.

Schindler, A. J., L. R. Baugh, and D. R. Sherwood. 2014. ‘Identification of late larval stage developmental checkpoints in Caenorhabditis elegans regulated by insulin/IGF and steroid hormone signaling pathways’, PLoS Genet, 10: e1004426.

Schulenburg, H., and M. A. Felix. 2017. ‘The Natural Biotic Environment of Caenorhabditis elegans’, Genetics, 206: 55–86.

Seidel, H. S., and J. Kimble. 2011. ‘The oogenic germline starvation response in C. elegans’, PLoS One, 6: e28074.

Seidel, H. S., M. V. Rockman, and L. Kruglyak. 2008. ‘Widespread genetic incompatibility in C. elegans maintained by balancing selection’, Science, 319: 589–94.

Srivastava, A., A. P. Morgan, M. L. Najarian, V. K. Sarsani, J. S. Sigmon, J. R. Shorter, A. Kashfeen, R. C. McMullan, L. H. Williams, P. Giusti-Rodriguez, M. T. Ferris, P. Sullivan, P. Hock, D. R. Miller, T. A. Bell, L. McMillan, G. A. Churchill, and F. P. de Villena. 2017. ‘Genomes of the Mouse Collaborative Cross’, Genetics, 206: 537–56.

Viney, M., and A. Diaz. 2012. ‘Phenotypic plasticity in nematodes: Evolutionary and ecological significance’, Worm, 1: 98–106.

Weigel, D. 2012. ‘Natural variation in Arabidopsis: from molecular genetics to ecological genomics’, Plant Physiol, 158: 2–22.

